# Identifying Mechanisms of Regulation to Model Carbon Flux During Heat Stress And Generate Testable Hypotheses

**DOI:** 10.1101/340950

**Authors:** Allen H. Hubbard, Xiaoke Zhang, Sara Jastrebski, Susan J. Lamont, Abhyudai Singh, Carl Schmidt

## Abstract

Understanding biological response to stimuli requires identifying mechanisms that coordinate changes across pathways. One of the promises of multi-omics studies is achieving this level of insight by simultaneously identifying different levels of regulation. However, computational approaches to integrate multiple types of data are lacking. An effective systems biology approach would be one that uses statistical methods to detect signatures of relevant network motifs and then builds metabolic circuits from these components to model shifting regulatory dynamics. For example, transcriptome and metabolome data complement one another in terms of their ability to describe shifts in physiology. Here, we extend a previously described method used to identify single nucleotide polymorphism (SNP’s) associated with metabolic changes (Gieger *et al.,* 2008). We apply this strategy to link changes in sulfur, amino acid and lipid production under heat stress by relating ratios of compounds to potential precursors and regulators. This approach provides integration of multi-omics data to link previously described, discrete units of regulation into functional pathways and hypothesizes novel biology relevant to the heat stress response.

## Introduction

The highly conserved heat stress response has been extensively studied in organisms across taxa. Understanding how to mitigate effects of hyperthermia has important applications in a variety of disciplines. For example, heat stroke is a common, severe complication of acute hyperthermia and a stronger biological understanding of its underpinnings could lead to more effective therapies (Kovats and Hajat, 2007). In the agricultural setting, prolonged heat stress - such as that encountered by livestock during heat waves - can decrease feed efficiency and animal growth and cause significant commercial losses in meat production (Lara *et al.,* 2013). Additionally, heat stress has been known to exert systemic physiological consequences such as changes in egg production and immune cell counts (Mashaly *et al.,* 2004).

While many of physiological consequences of hyperthermia are understood, a strong mechanistic description is lacking.

Metabolic shifts during heat shift involve a number of signaling cascades (Verghese *et al.,* 2012). These include the unfolded protein response and both pro and anti-apoptotic pathways (Fulda *et al.,* 2010). Increasing evidence suggests that biologically active lipids play an important function during heat stress, as signaling agents and maintaining cell membrane integrity (Balogh *et al.,* 2013). Establishing the relationship between lipid metabolism and other well-characterized heat responsive pathways, would provide better understanding of the flow of resources to different types of metabolites. This could provide a model describing how small carbon precursors are selectively routed to various fates necessary to sustain signaling, energy production and other processes that must undergo dynamic shifts under heat stress.

Viewing carbon flow as a circuit with dynamics that are affected by gene expression changes produces an effective description and identifies testable biological hypotheses. This perspective describes the mechanisms that manage and create resource pools in the form of biochemically valuable carbon backbones, including cysteine and other catabolized amino acids that are selectively incorporated into various biologically active of molecules. The model that we construct describes the interconnection between production of antioxidants, gluconeogenesis, and production of signaling and structural lipids.

Redirection of carbon backbones occurs at specific points of regulation where molecules are processed into one of two or more available metabolic fates. We have previously introduced this type of regulation in the form of metabolic forks, but have now extended them to build pathway level models (Hubbard *et al.,* 2018). Importantly, this arrangement allows gene expression patterns to implement regulatory logic by selectively directing resources.

## Materials and Methods

We first subsetted metabolite measurements for compounds representing sulfur, lipid and sugar metabolism because these are major processes influenced by heat stress (Jastrebski *et al.,* 2017). This initial step of feature selection reduced metabolite data from 600 compounds to approximately 60. Our analysis deliberately focused on regulatory circuits controlling metabolism involving only the selected components of tissue enriched genes and metabolites linking sulfur and lipid metabolism.

**Figure 1:**
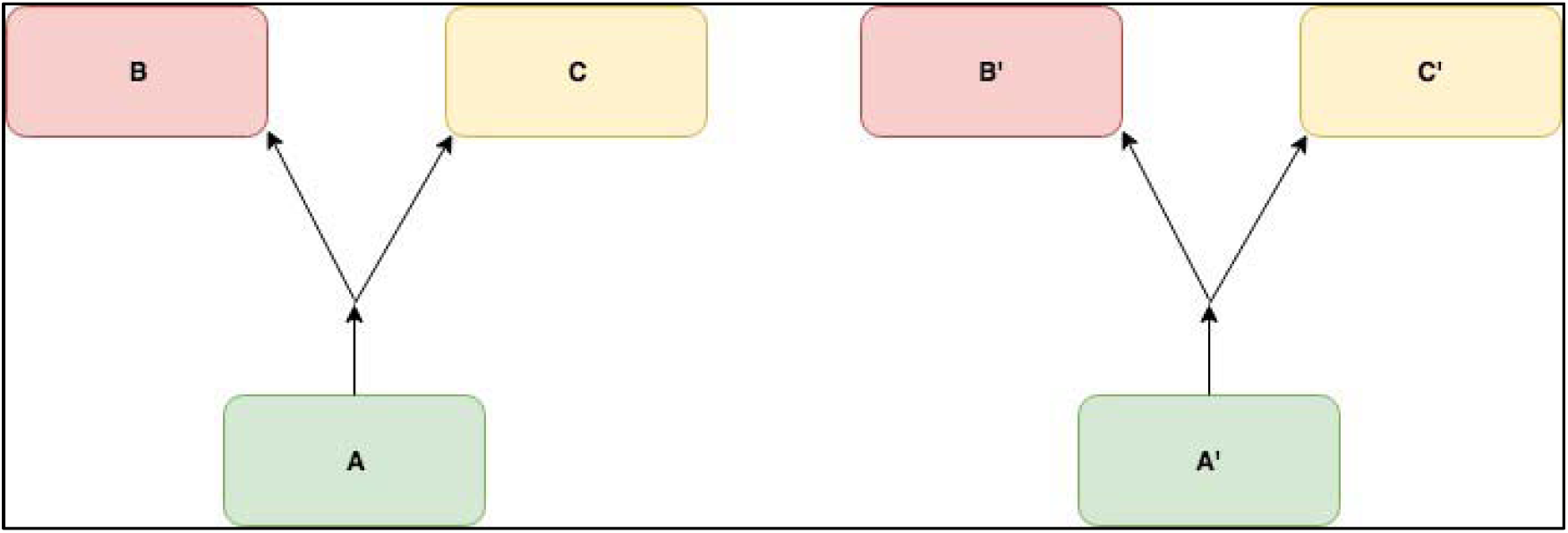
An example of two potential metabolic forks detected from the data through linear models. Each one represents a triplet whose models 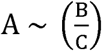 and 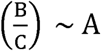 have significant interaction terms between control and heat stress conditions.

**Figure 2:**
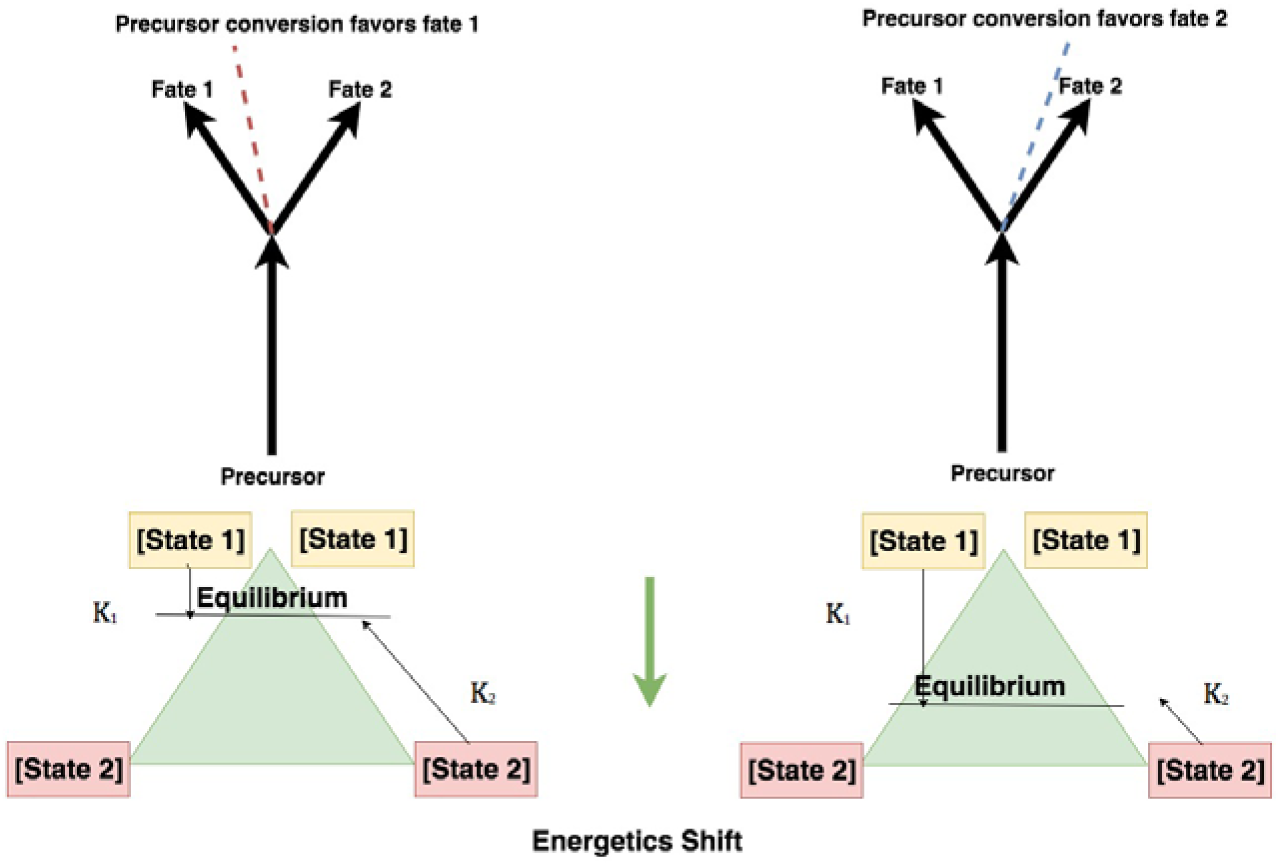
Metabolic forks, representing different favorabilities for the metabolic fates at regulatory branch points. A shift in regulation changes the energetic favorability in favor of one route of the metabolic fork.

**Figure 2:**
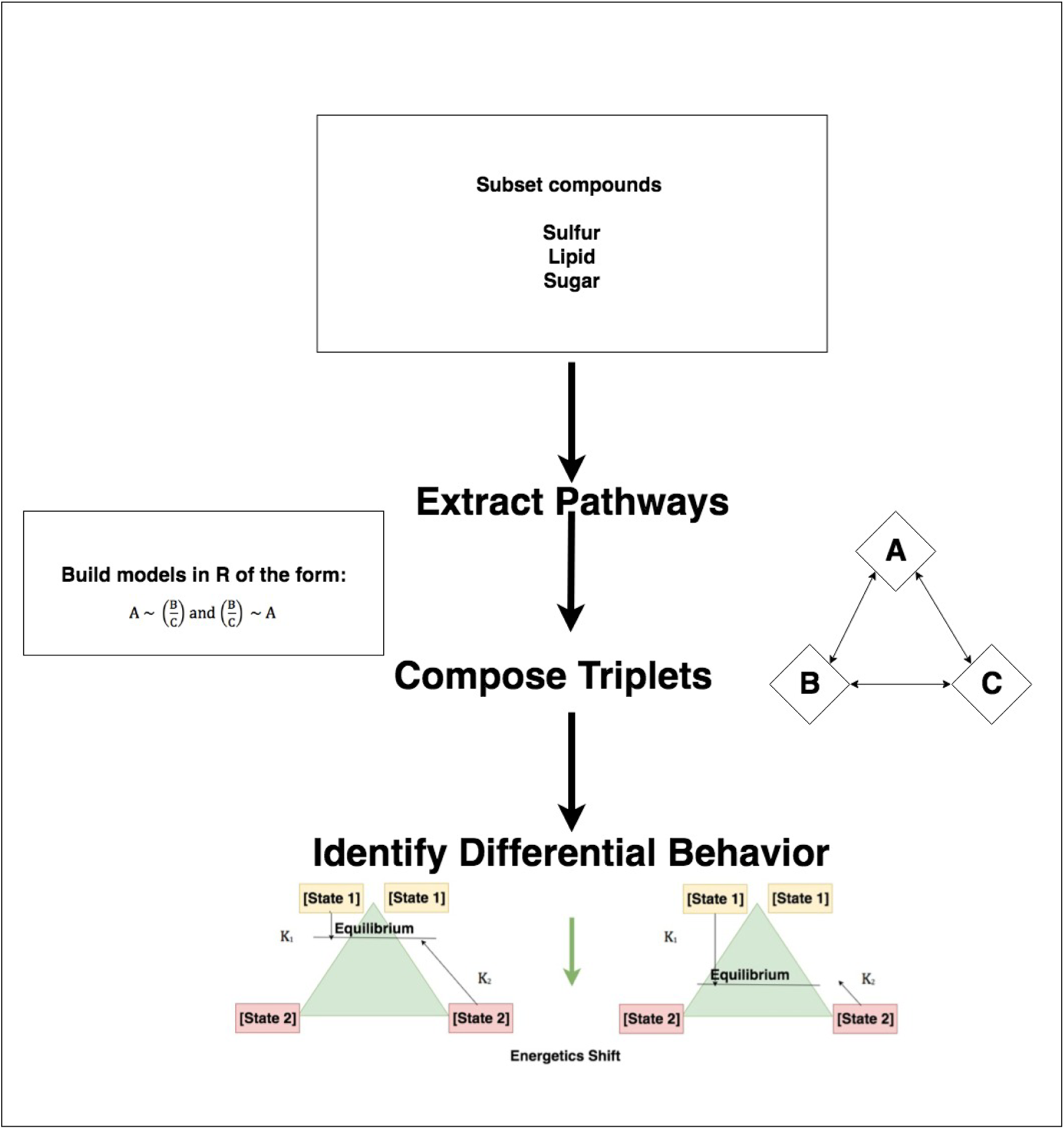
Workflow to identify triplets of compound that regulate sulfur and lipid metabolism.

### 5.1 Iterative Linear Models And Metabolic Forks

We evaluated levels of compounds representing sulfur, lipid and sugar metabolism as these are major processes influenced by heat stress, in order to determine metabolic forks among these systems. Then, values for the correlation function of the form 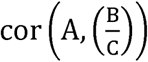 were calculated where A,B and C represents the levels of metabolites or gene transcripts. The correlation between compound A and the ratio 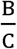 can differ between control and experimental conditions when there is heat responsive regulation of members of the triplet. The most biologically informative triplets of the form A, B and C often represent sets of precursors and their resulting metabolic products. Ratios of compounds are used in these functions because they are more sensitive to detecting points of potential regulation for diverging metabolic routes (Gieger *et al.,* 2008). A biochemical interpretation of these functions is provided in Figure 2. Triplets whose difference in value for the correlation function was 1.2 or greater between control and experimental conditions, i.e. 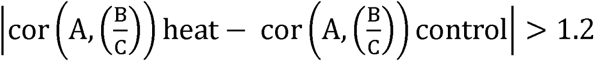, were selected as representing possible metabolic forks. A threshold was chosen because of the trade-off between identifying a diverse set of triplets and the need to achieve stringency such that resulting linear models of the triplet were likely to have significant interaction terms. Linear models were then used to detect differential behavior under heat stress and to identify triplets with significant interaction terms – indicating different slopes between control and heat stress conditions. The threshold of 1.2 for identifying potentially interesting triplets was ad-hoc, with the linear models being used to determine a p-value for the difference between control and experimental conditions. Thus, ad-hoc correlation functions were used to propose candidates for more rigorous evaluation by linear models. To be considered as a pathway element and incorporated into a circuit, the interaction term must be significant for both models of the form 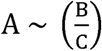 and 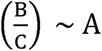. This stringent heuristic is chosen because of ambiguity regarding directionality of the relationship between 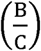 and A. For example, while ratios are more sensitive to detecting relationships between possible sets of precursors and a product, it is not always clear which relationships among the triplet are causal and which are correlative. While the reliance on linear models does enforce an assumption of linearity such an assumption is in accordance with the use of correlation, which also measures linear relationships, to identify differential regulation of triplets. Triplets were merged with one another to generate pathways. We focus on three involving sulfur and lipid regulation, because we believe that the regulation associated with these triplets represents the functioning pathway of lipid and antioxidant regulation also described by transcriptome data. Regarding components of a metabolic fork, in terms of their relationship to one another as precursors and products, these hypotheses are necessarily associative and not always causal. However, confidence in the proposed directionality of relationships can be strengthened by gene expression changes. Per existing methods, all data was log transformed before modeling (Illig *et al,* 2010). Once libraries were sequenced, data were processed using an in-house pipeline and fragments per kilobase per million mapped reads (FPKM) values were determined. Differential expression was determined by using the standard t.test function in R.

**Figure 3:**
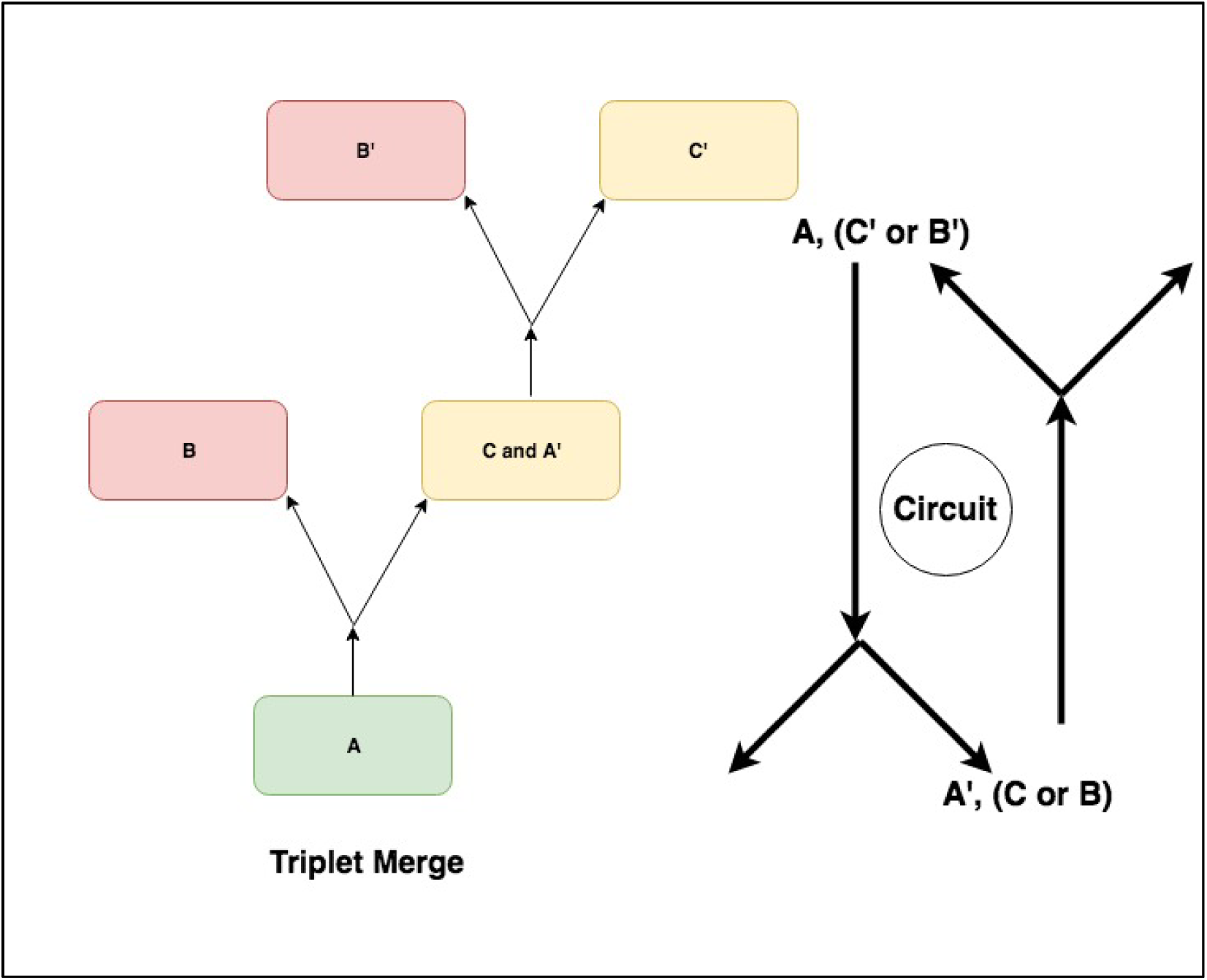
Metabolic forks being merged. Metabolic forks are joined into potential pathways by identifying forks that share overlapping members of triplets.

By merging together forks it was possible to identify small, functional units that may be critical elements of pathways. Integrating these isolated units to form controlled regulatory systems can identify circuits of carbon and sulfur regulation. Importantly, linear models relying on the ratios of metabolites identify differential behavior not detectable using raw expression measurements alone (Fig. 2). This may be due to reductions in variance (Gieger *et al,* 2008) as well as an ability to capture underlying biology by being more sensitive to fluxes down each metabolic pathway. These models were joined to create a larger circuit of regulation, in which each metabolic fork contains a component that is also a member of other metabolic forks.

### From Merged Triplets to Pathways

By merging together forks it is possible to identify small, functional units that may be critical elements of pathways. Integrating these isolated units to form controlled regulatory systems can identify circuits of carbon and sulfur regulation. Importantly, linear models relying on the ratios of metabolites identify differential behavior not detectable using raw expression measurements alone (Fig. 2). This may be due to reductions in variance (Gieger *et al,* 2008) as well as an ability to capture underlying biology by being more sensitive to fluxes down each metabolic pathway. Importantly, these models can then be joined to create a larger circuit of regulation. In these types of models, elements involved in each metabolic fork also play a role in the functioning of other metabolic forks. The metabolic interpretation of each metabolic fork, and the joining of multiple examples to form circuits can capture the intuition and biochemistry of pathways.

## Results And Discussions

One such circuit, predicted from the data, relates involving lipid and anti-oxidant compounds (Fig. 3). These predictions are consistent with previous research relating hypercysteinemia and hyperlipidemia to one another (Herman and Obeid, 2009).

**Fig. 2:**
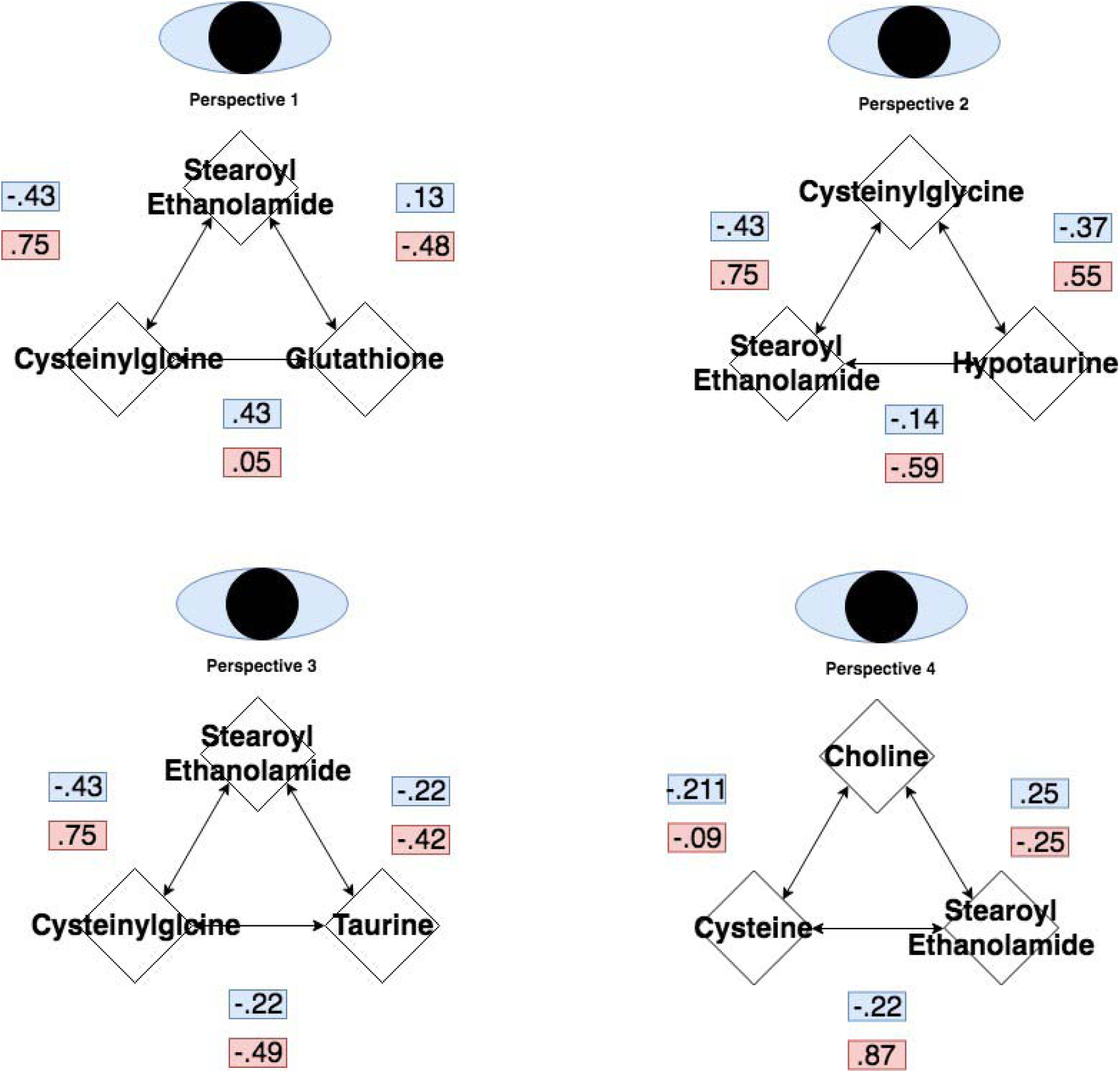
Metabolic forks joined together describe a pathway that regulated by changes in gene regulation.

**Fig. 3:**
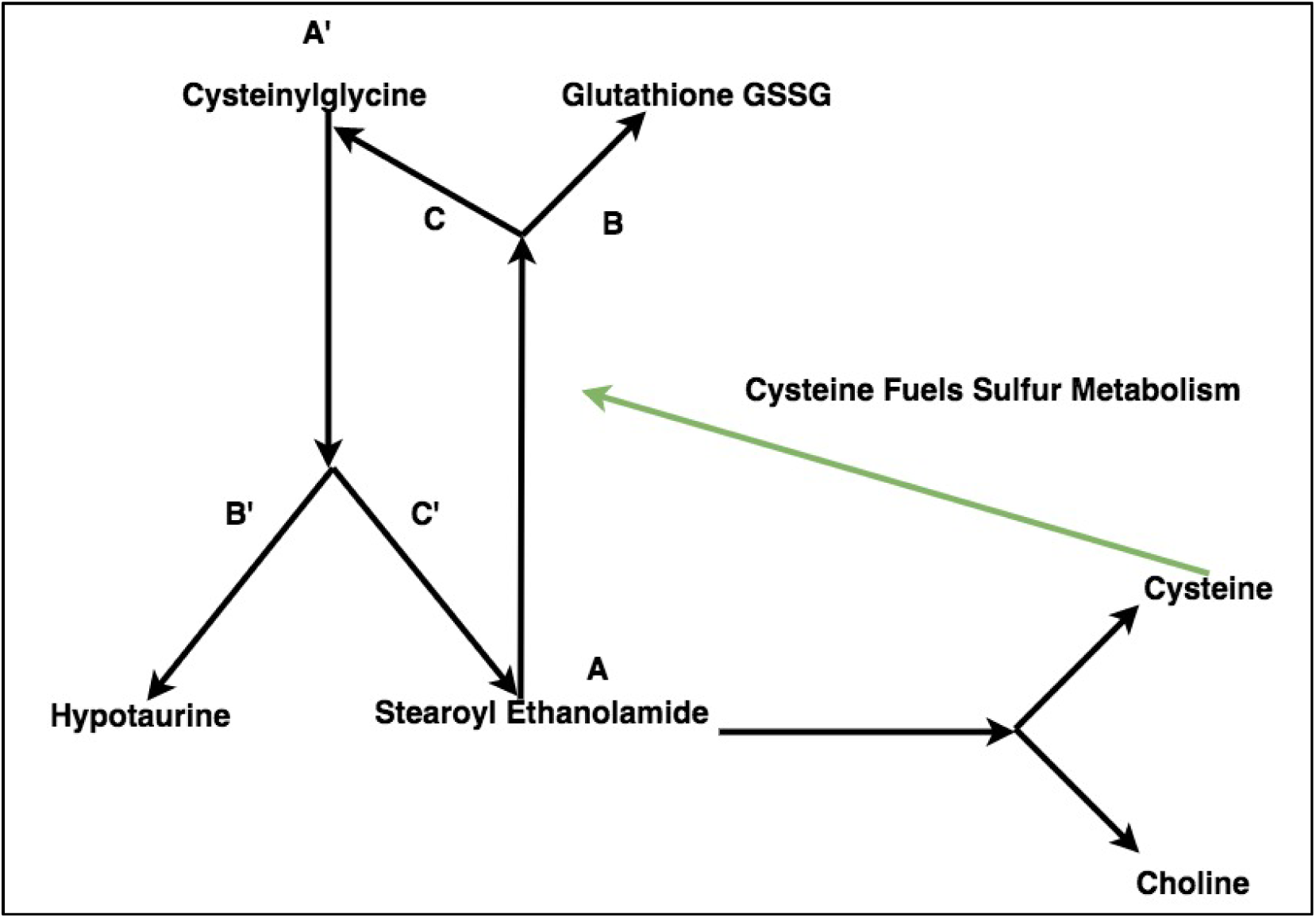
Extended circuit based off of merging of triplets

A full mechanism relating sulfur, lipid and antioxidant activities to one another can be constructed by joining additional triplets (Fig. 3). This is done by joining triplets that share at least one overlapping element. In this model, cysteine levels are increased under heat stress (Jastrebski *et al.,* 2017), driving sulfur metabolism coupled to lipid and antioxidant production via changes in expression of key regulatory genes. Redirection of resources to cysteine metabolism is hypothesized to occur at the expense of choline derived signaling and structural lipids, due to changes in gene expression for enzymes related to this process. Differential behavior at each of these forks can be seen in a series of linear models, each of which demonstrate significant interaction terms (Fig. 4). These branch points are then placed in context of known biology to generate a regulatory model that incorporates both transcriptome and metabolome measurements. Genes that regulate the metabolism of each metabolite in the network skeleton can be located on the skeleton in order to flesh out a full pathway model.

**Fig. 4A-F.**
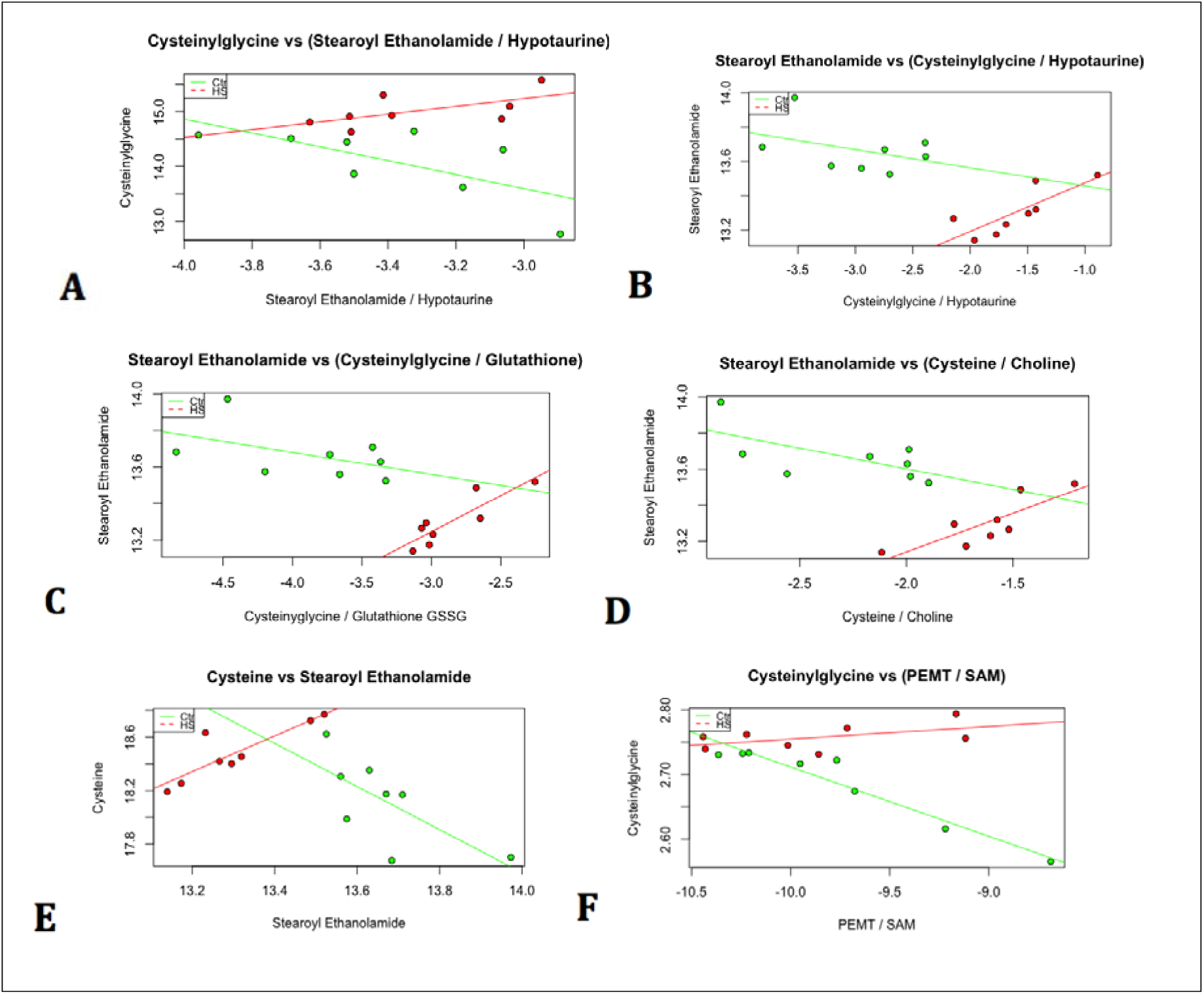
**Metabolic Forks and Related Models:** Linear models detect differential behavior of the metabolic forks that comprise the circuit. Figures (6-9) describe each branch-point in detail. All p-values for relevant interaction terms are less than .05. Data points that are green represent measurements from control conditions, whereas red data points represent measurements from heat stressed birds. Discussion

Fig. 5 is a proposed metabolic circuit relating lipid, cysteine and glutathione production. It contains sets of metabolic forks at key regulatory points, operating in concert. Many of these regulate the incorporation of sulfur into biologically important molecules. For example, under heat stress, sulfur metabolism favors glutathione at the expense of taurine (Fig. 6). This is controlled by a metabolic fork, which is critical to antioxidant production (Fig. 7). Cysteine, which fuels many sulfur processing pathways, is the only amino acid increased under chronic heat stress. Our work describes mechanisms by which pools of cysteine regulate the long-term heat stress response. This model contextualizes mechanisms predicted by metabolite data with transcriptome data and known biology.

**Fig. 5:**
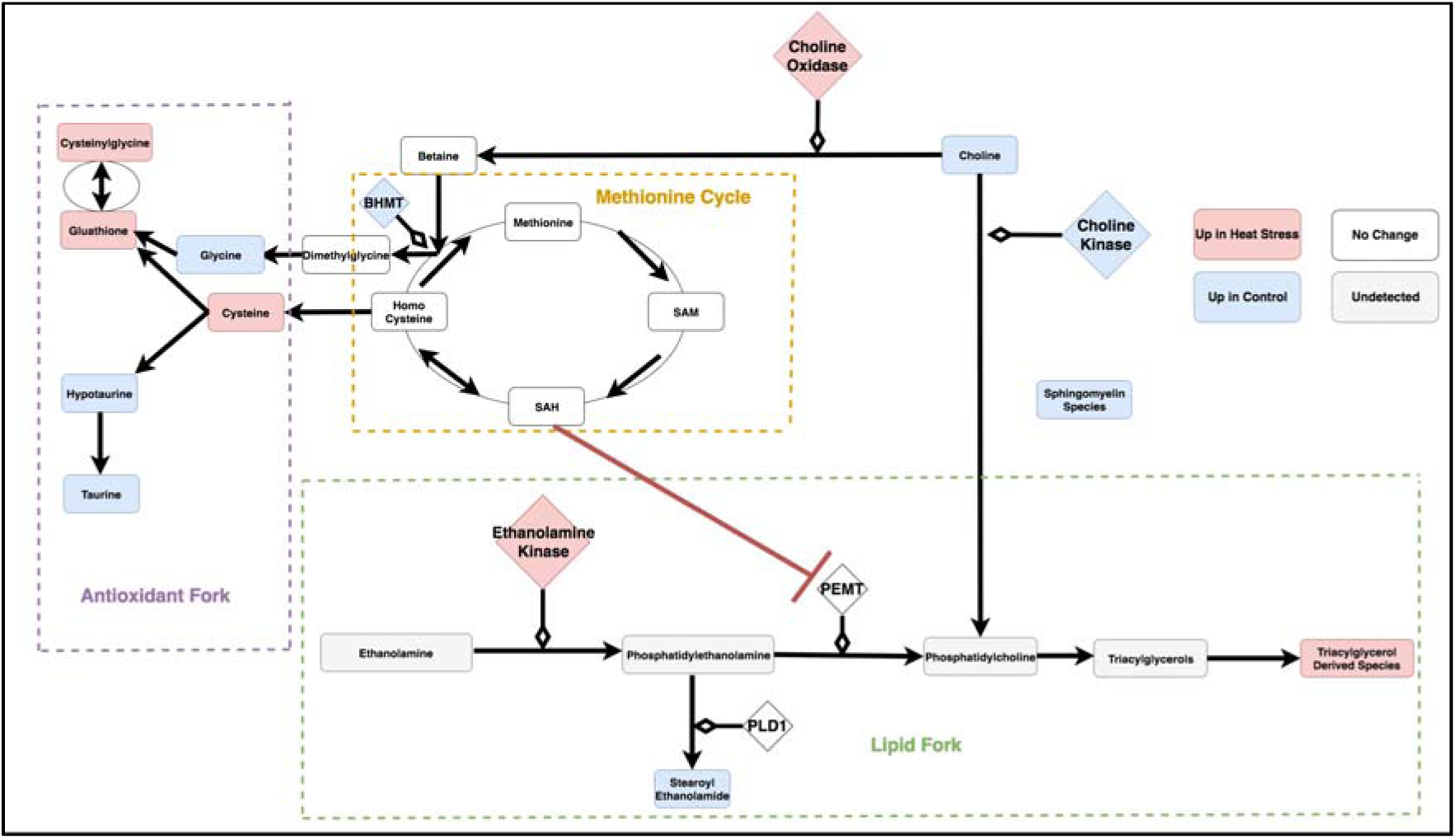
The circuit components as modules summarized by the three categories of antioxidant, lipid and methionine metabolism. SAM: S-Adenosyl-L-methionine, SAH: S-Adenosyl-L-homocysteine, Glutathione GSSG: Glutathione Disulfide, PEMT: Phosphatidylethanolamine N-methyltransferase, BHMT: Betaine--Homocysteine S-Methyltransferase, PLD1: Phospholipase D-1, PEMT: Phosphatidylethanolamine N-methyltransferase.

Changes in expression among genes regulating the methionine cycle are critical to the activity of this pathway. Additionally, S-Adenosyl-L-homocysteine (SAH) and Phosphatidylethanolamine N-methyltransferase (PEMT) interactions are linked to ethanolamide metabolism in this model. A similar relationship between methionine metabolism and PEMT features in a putative relationship between hyperlipidemia and hyperhomocysteinemia (Obeid and Hermann, 2009). Changes to sulfur metabolism under heat stress influence lipids in a number of ways beyond the SAH interaction with PEMT. In our model, choline, the precursor to many fatty acids, is directed away from the production of signaling and structural lipids. Several shifts in gene expression route this resource towards sulfur metabolism. Choline oxidase, the gene encoding the enzyme oxidizing choline to produce betaine, is also up-regulated. Betaine plays an important role in the methionine cycle by providing an alternative pathway for methylation of homocysteine (Kempson *et al.,* 2014). Concurrently, transcription of the first enzyme involved in converting choline to phosphatidylcholine, choline kinase, is down-regulated. Betaine levels, however, are unchanged, suggesting redirected choline rescues betaine levels. Further supporting a relationship between sulfur and lipid metabolism via choline and betaine, Betaine--Homocysteine S-Methyltransferase (BHMT) transcription is downregulate under heat stress. BHMT converts betaine and homocysteine to dimethylglycine and methionine, and mouse knockouts of this gene show highly elevated levels of homocysteine (Strakova *et al.,* 2012).

Phosphoethanolamine kinase is up-regulated, consistent with changes to prevent depletion of phosphatidylethanolamine-derived lipids such as phosphatidylcholine and stearoyl ethanolamide. This model predicts that maintaining phosphatidylcholine production, despite dramatically directing resources to antioxidant production may be critical to homeostasis.

### Regulation of Individual Forks

Ultimately, this model involving selective processing of sulfur increases the reservoir of anti-oxidants. The circuit also proposes changes in lipids, including coupling ethanolamine related compounds, such as phosphatidylethanolamine and stearoyl ethanolamide, is coupled to glutathione production through cysteine processing pathways. In our model, this is accomplished through changes in the methionine cycle and choline metabolism. These mechanisms are consistent with gene expression changes. Under heat stress, decreased BHMT transcription preserves cysteine pools, managing the activity of the methionine to S-Adenosyl-L-methionine (SAM)/S-Adenosyl-L-homocysteine (SAH) cycle. A major product of this cycle, SAH, is a potent inhibitor of Phosphatidylethanolamine N-methyltransferase (PEMT). Because higher levels of cysteine would ordinarily fuel methionine metabolism, the balance between cysteine allocation to antioxidants and the methionine cycle may be a critical for lipid metabolism.

**Fig. 6:**
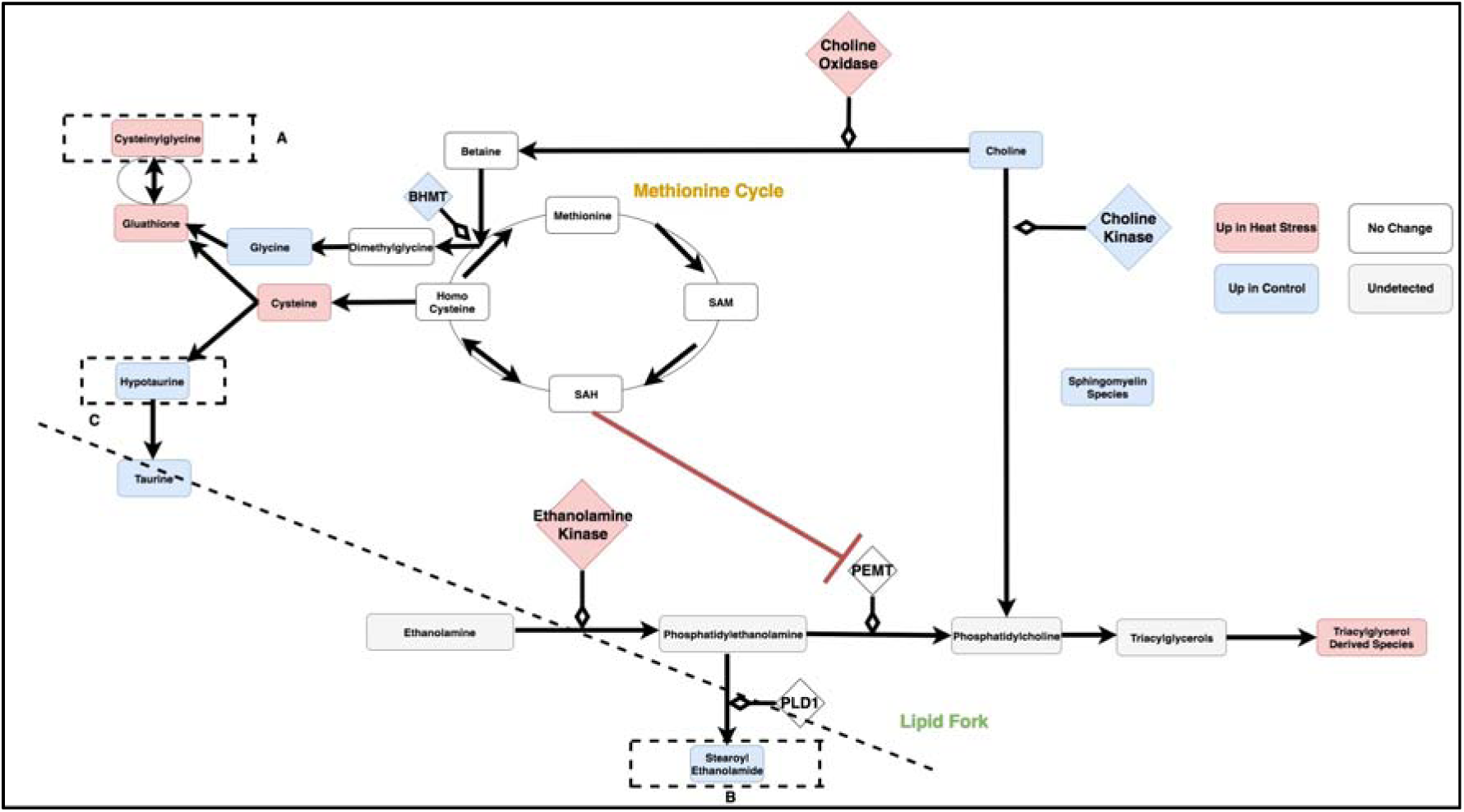
Triplet of cysteinylglycine and (stearoyl ethanolamide / hypotaurine). The compartmentalization of the pathway by regions containing the compounds in the ratio (stearoyl ethanolamide and hypotaurine) is illustrated by the dotted line. For the linear model representing differential behavior of this branch point, see Fig. 4A. SAM: S-Adenosyl-L-methionine, SAH: S-Adenosyl-L-homocysteine,Glutathione GSSG: Glutathione Disulfide, PEMT: Phosphatidylethanolamine N-methyltransferase, BHMT: Betaine--Homocysteine S-Methyltransferase, PLD1: Phospholipase D-1, PEMT: Phosphatidylethanolamine N-methyltransferase.

Under heat stress conditions, stearoyl ethanolamide levels correlate well with ratios of the reduced glutathione derivative, cysteinylglcyine, and taurine (Fig. 7). This latter quantity represents a metabolic fork underlying sulfur metabolism, which favors glutathione under heat stress. Under control conditions, activation of the sulfur metabolism would inhibit an important component of stearoyl ethanolamide production via SAH-related inhibition of PEMT. This mechanism is countered under heat stress conditions with an increase in the ratio of PEMT/SAM correlating with rising levels of gamma glutamylcysteine (Fig. 4F).

**Fig. 7:**
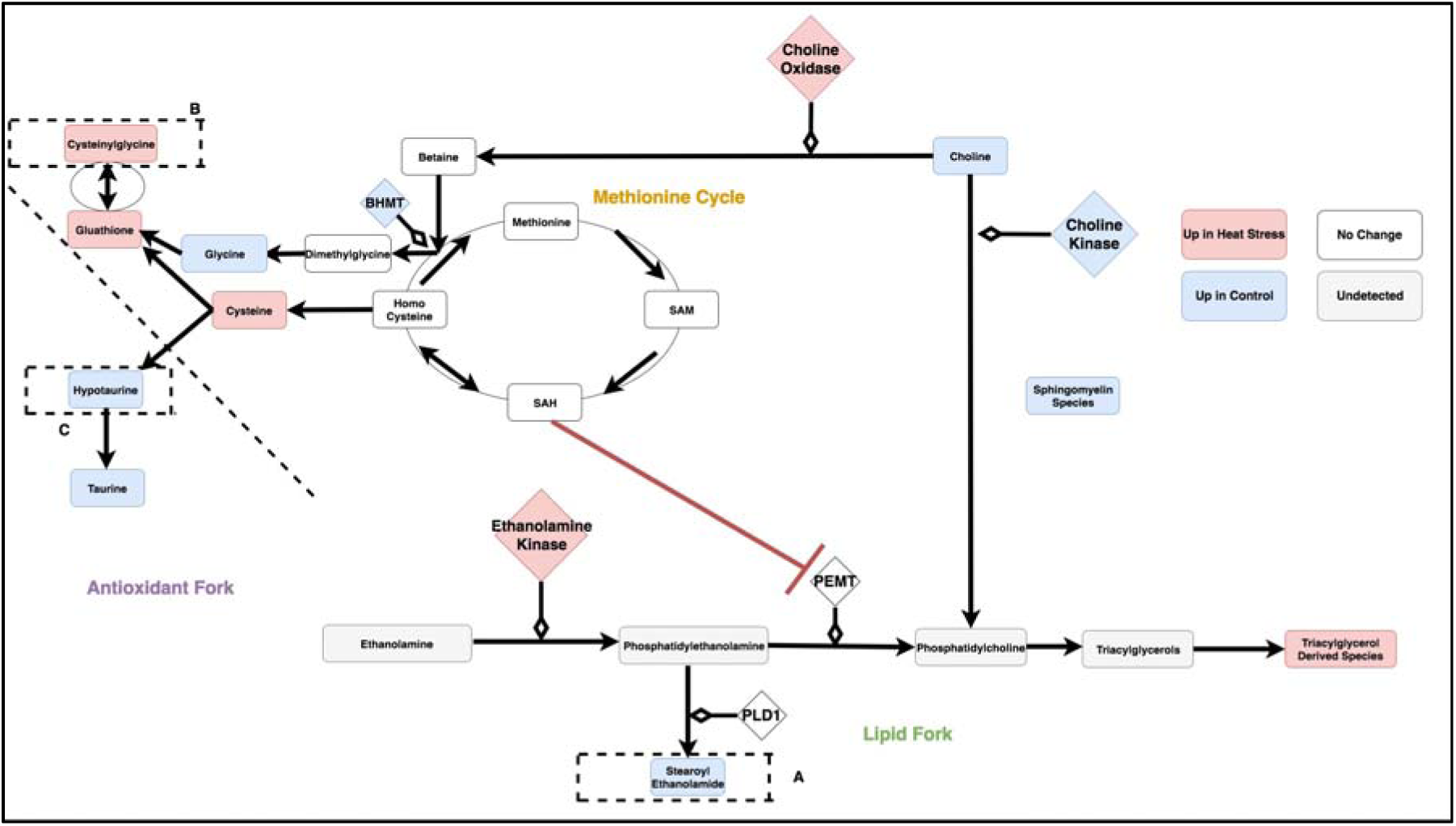
Triplet of stearoyl ethanolamide and (cysteinylglycine / hypotaurine). The compartmentalization of the pathway by regions containing the compounds in the ratio (cysteinylglycine and hypotaurine) is illustrated by the dotted line. For the linear model representing differential behavior of this branch point, see Fig. 4B. SAM: S-Adenosyl-L-methionine, SAH: S-Adenosyl-L-homocysteine,Glutathione GSSG: Glutathione Disulfide, PEMT: Phosphatidylethanolamine N-methyltransferase, BHMT: Betaine--Homocysteine S-Methyltransferase, PLD1: Phospholipase D-1, PEMT: Phosphatidylethanolamine N-methyltransferase.

Stearoyl ethanolamide levels and the ratio of the reduced glutathione derivative, cysteinylglycine, and Glutathione GSSG (Glutathione Disulfide) show strong patterns of differential correlation between control and heat stress (Fig. 8). This is consistent with concerted regulation of several metabolic forks in the underlying circuit of carbon metabolism.

**Fig. 8:**
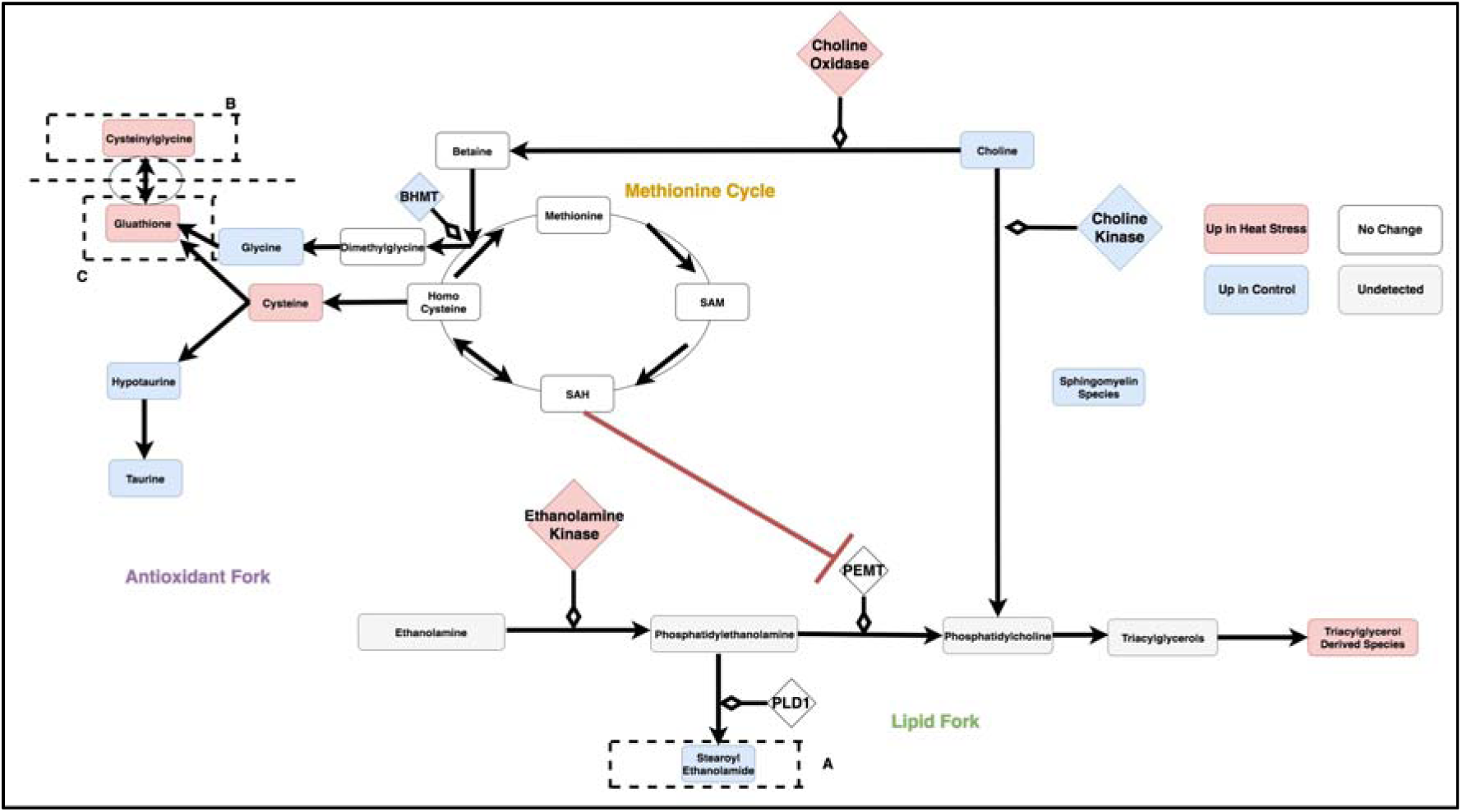
Triplet of stearoyl ethanolamide and (cysteinylglycine / gluathatione). The compartmentalization of the pathway by regions containing the compounds in the ratio (cysteinylglycine and glutathione) is illustrated by the dotted line. For the linear model representing differential behavior of this branch point, see Fig. 4C. SAM: S-Adenosyl-L-methionine, SAH: S-Adenosyl-L-homocysteine,Glutathione GSSG: Glutathione Disulfide, PEMT: Phosphatidylethanolamine N-methyltransferase, BHMT: Betaine--Homocysteine S-Methyltransferase, PLD1: Phospholipase D-1, PEMT: Phosphatidylethanolamine N-methyltransferase.

Stearoyl ethanolamide levels and the ratio of cysteine and choline shows strong patterns of differential correlation between control and heat stress (Fig. 9). Under the proposed mechanism, as cysteine metabolism is increased during heat stress, choline decreases with its remaining levels critical to maintain betaine.

**Fig. 9:**
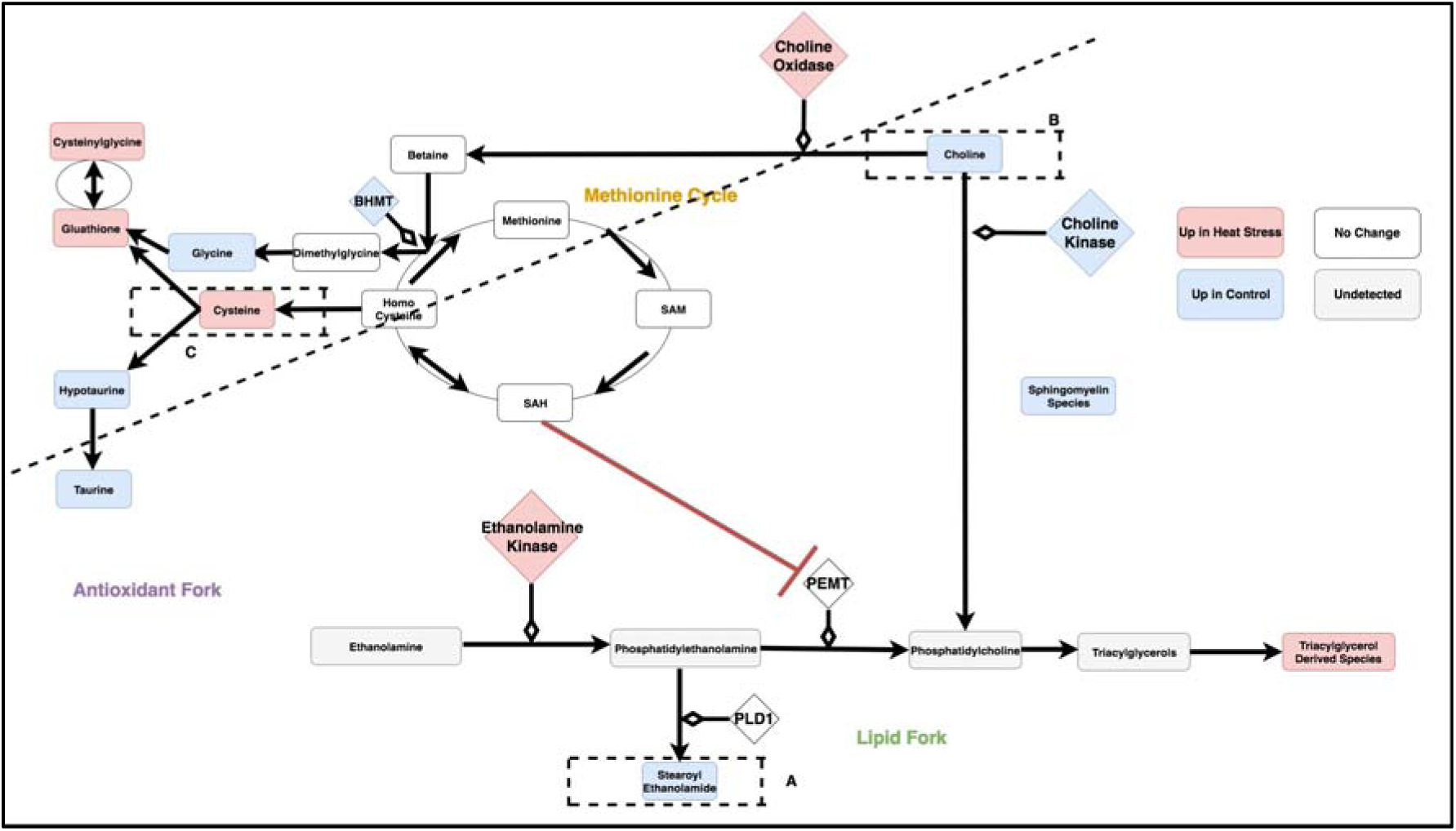
Triplet of stearoyl ethanolamide and (cysteine / choline). The compartmentalization of the pathway by regions containing the compounds in the ratio (choline and cysteine) is illustrated by the dotted line. For the linear model representing differential behavior of this branch point, see Fig. 4D. SAM: S-Adenosyl-L-methionine, SAH: S-Adenosyl-L-homocysteine, Glutathione GSSG: Glutathione Disulfide, PEMT: Phosphatidylethanolamine N-methyltransferase, BHMT: Betaine--Homocysteine S-Methyltransferase, PLD1: Phospholipase D-1, PEMT: Phosphatidylethanolamine N-methyltransferase.

## Conclusions

This circuit driven model, derived by joining metabolic forks as network elements, provides a mechanistic context for correlations that would otherwise be enigmatic. For example, stearoyl ethanolamide and cysteine demonstrate differential relationships between control and heat stress conditions (p-value of interaction term < .05). The physiological roles of stearoyl ethanolamide are not fully established, although it has been shown to have anti-inflammatory properties (Ezzili *et al.,* 2010). This makes the compound similar to many other metabolites involved in the heat stress response. Though stearoyl ethanolamide is downregulated, its correlation with cysteine indicates regulatory coupling during heat stress response. This connection would support a metabolic circuit connecting antioxidant and lipid metabolism. Transcriptome data also supports increased utilization of pathways for cysteine production under heat stress in a way that influences lipid production (Fig. 4E). This combination of transcriptome and metabolome data in order to achieve biological insight, helps further a critical goal of modern genomics: to determine the mechanisms that control physiology. Our computational analysis provides insight into changes that may influence physiology under heat stress. This analysis provides network context for previous studies that have identified differential gene expression and metabolite levels (Jastrebski *et al.,* 2017). Systems biology studies can use multi-omics data to identify elements of regulation, integrating them into concrete networks that generate hypotheses about large-scale regulation. Collectively, these changes in gene expression and metabolic forks identified by this work provide mechanistic context for the differential relationship between stearoyl ethanolamide and cysteine during heat stress. The insights from this study expand the role of carbon of and sulfur flux during the long-term heat stress response.

This work provides clarification of previous studies that explore feed supplementation, albeit with incomplete models of the circuit currently proposed. For example, betaine and choline supplementation has variable effects on bird performance with recent studies suggesting it has limited influence on improving broiler performance and cannot overcome the negative influences of heat stress (Kpodo *et al.,* 2015). According our extended circuit, the bird is able to effectively maintain betaine levels under heat stress through redirection of choline such that supplementation may be ineffective at shifting network dynamics. We hypothesize gene regulation shunts choline to betaine, preventing the accumulation of resource deficits. Such changes include downregulation of BHMT, upregulation of choline oxidase and downregulation of choline kinase. The impact of these changes could be dramatic. Modification of choline accounts for 70 percent of phospohatidylcholine synthesis, with the remaining 30 percent derived from PEMT driven methylation of phosphotidylethanolamine (DeLong *et al.,* 1999). This latter pathway also has gene expression changes, such as up-regulation of choline kinase. These changes may compensate for altered choline dynamics during long-term heat stress. Thus, the ability of choline to rescue performance from stress may be stress and organism specific. For example, choline supplementation has been shown in clinical studies to improve antioxidant efficiency in cystic fibrosis patients (Innis *et al.,* 2007) despite its efficacy in influencing livestock performance being equivocal (Kpodo *et al.,* 2015).

The hypotheses generated by this work propose mechanisms that may underlie associations from GWAS (genome wide association studies). This is important, as previous work on quantifying broiler performance under heat stress, has relied on QTL (quantitative trait loci) mapping to identify potentially relevant SNP’s controlling relevant metrics. One of these resides in the PEMT gene, implicated in sulfur and lipid metabolism, as being associated with body temperature at Day 20 (Van Goor *et al,* 2015). Our proposed circuit includes PEMT as a critical element in a broader network and provides a possible functional role of the previously identified SNP. Building circuits from individual network units provides biological context for statistical observations in a way that relate components from different, but connected, pathways.

Our approach ultimately creates a network integrating compounds whose role in heat stress are well understood and compounds not previously implicated in the heat stress response. This is particularly useful for providing insight into the relatively uncharacterized lipid, stearoyl ethanolamide, that is altered by heat stress. Stearoyl ethanaolamide is as an example of an n-acylamide endocannabinoid. The functions of n-acylamides are best understood in the brain, where they play important roles in signaling and inflammation response (Raboune *et al.,* 2014). Thus, changes in ratios of these species can be informative as they may represent preferential routing of carbon resources to lipids that influence inflammation.

The shifts in compounds such as stearoyl ethanolamide are also consistent with a circuit preferentially directing carbon backbones towards gluconeogenesis and triacylglycerol production. Under this complete model, the bird allocates carbon resources to signaling molecules as well as to antioxidant and energy production pathways. Leveraging computational methods to understand the nuances of carbon and sulfur flow under heat stress provides a significant improvement in understanding the regulation of the response, and generates a number of testable hypotheses. These are being incorporated to plan studies in which feed composition is altered with resources thought to be involved in the major circuits. Additionally, we have successfully captured the logic of the carbon flow under heat stress. The transition from simply determining up or down regulation of certain compounds developing a collection of well-characterized mechanisms to be integrated into circuits is a powerful improvement in using systems biology to integrate large-scale multi-omics data.

## Availability of Data and Materials

Transcriptome sequencing data is publicly available through GEO series accession number GSE95088 (https://www.ncbi.nlm.nih.gov/geo/query/acc.cgi?acc=GSE95088). <u>Metabolome data is included as supplementary data</u>.

## Acknowledgements

We thank Greg Keane and members of the University of Delaware IT staff for their valuable assistance, and additional members of the Schmidt group for valuable comments.

## Abbreviations

SNP: Single Nucleotide Polymorphism
Glutathione GSSG: Glutathione Disulfide
SAH: S-Adenosyl-L-homocysteine
SAM: S-Adenosyl-L-methionine
PEMT: Phosphatidylethanolamine N-methyltransferase
BHMT: Betaine--Homocysteine S-Methyltransferase
PLD-1: Phospholipase-D1
GWAS: Genome Wide Association Studies
QTL: Quantitative Trait Loci

